# The sialidase NEU3 promotes pulmonary fibrosis in mice

**DOI:** 10.1101/2022.02.12.480197

**Authors:** Darrell Pilling, Kyle Sahlberg, Tejas R. Karhadkar, Wensheng Chen, Richard H. Gomer

## Abstract

Sialic acid is often the distal sugar on glycoconjugates, and sialidases are enzymes that remove this sugar. In fibrotic lesions in human and mouse lungs, there is extensive desialylation of glycoconjugates, and upregulation of sialidases including the extracellular sialidase NEU3. In the bleomycin model of pulmonary fibrosis, mice lacking NEU3 (*Neu3*^*-/-*^) showed strongly attenuated bleomycin-induced weight loss, lung damage, inflammation, upregulation of TGF-β1, and fibrosis. This indicates that NEU3 is necessary for the full spectrum of bleomycin-induced pulmonary fibrosis. To determine if NEU3 is sufficient for fibrosis, mice not treated with bleomycin were treated with recombinant murine NEU3 or inactive NEU3. Aspiration of NEU3 caused inflammation and fibrosis in the lungs, while inactive NEU3 caused inflammation but not fibrosis. Mice were also treated with NEU3 starting 10 days after bleomycin. In male but not female mice, NEU3 increased inflammation and fibrosis. Inactive NEU3 did not enhance bleomycin-induced lung fibrosis. These results suggest that NEU3 is sufficient to induce fibrosis in the lungs, and that this effect is mediated by NEU3’s enzymic activity.

## Introduction

Fibrosing diseases, such as idiopathic pulmonary fibrosis (IPF), cardiac fibrosis, liver cirrhosis, and end-stage kidney disease, involve the progressive formation of scar tissue in internal organs that replaces the normal tissue, and are an increasing burden for healthcare systems [1-4]. IPF is a chronic and fatal disease that affects ∼3 million people worldwide, with an incidence of 1 in 200 > 65 years, and is characterized by fibrosis of the lungs and if untreated has a median survival of 3–5 years after initial diagnosis [5-7]. The only FDA-approved therapeutics, Nintedanib and Pirfenidone, slow but do not reverse the progression of the disease [6, 8].

Many secreted and cell-surface mammalian proteins are glycosylated, and many of the glycosylated structures have sialic acids as the monosaccharide at the end of the polysaccharide chain [9-11]. Sialidases, which are also called neuraminidases, remove the terminal sialic acid from these glycoconjugates [12, 13]. There are four known mammalian sialidases, NEU1, NEU2, NEU3, and NEU4, and each sialidase has different subcellular localizations and substrate specificities [12-14]. Of the four sialidases, only NEU3 is primarily extracellular [14-17]. This enzyme uses two tyrosines in the active site to remove sialic acid from glycoconjugates [13, 18].

The lungs from both preclinical models of lung fibrosis and patients with IPF have increased levels of sialidase activity in the bronchoalveolar lavage fluid (BALF), increased levels of NEU3 in the BAL, and increased levels of both NEU1 and NEU3 in the lung tissue [19-23]. Mice treated with NEU1 and NEU3 inhibitors, and *Neu3*^*-/-*^ knockout mice, show strongly attenuated inflammation and fibrosis after bleomycin challenge [22-26], suggesting that NEU3 is necessary for the full extent of bleomycin-induced inflammation and fibrosis.

In this report, we assessed whether NEU3 might be sufficient to induce fibrosis. We previously found that mice have low levels of NEU3 (∼1.7 ng) in their BALF, while mice treated with bleomycin have ∼15 ng of NEU3 in their BALF at day 21 [23]. We find that aspiration of 15 ng of mouse NEU3 every other day for 20 days caused inflammation and fibrosis in the lungs at day 21, and that the fibrosis was not observed using a mutated and thus inactive NEU3 lacking the two key tyrosines in the active site. We also find that NEU3 potentiates bleomycin-induced inflammation and pulmonary fibrosis. These results suggest that NEU3 is sufficient to induce fibrosis.

## Materials and methods

### Recombinant mouse NEU3 expression

Chinese hamster ovary (CHO-K1) cells were cultured in CDM4CHO medium with L-glutamine (Cat# SH30557.02, Hyclone/Cytiva, Marlborough, MA). Cells (1 × 10^5^) were mixed with 2 μg of 100 μg/ml of murine NEU3 expression plasmid (MR223297; Origene, Rockville, MD) in 100 μl PBS (GE Lifesciences, Marlborough, MA) and were transfected by electroporation using a 4D-Nucleofector System (Lonza, Walkersville, MD) following the manufacturer’s protocol. Before use, the plasmid was sequenced for verification as previously described [25, 27-29]. The transfected cells were kept at room temperature for 15 minutes for recovery, after which the CHO-K1 cells were cultured in 25 ml CDM4CHO medium in a humidified incubator at 37°C with 5% CO_2_. After 24 hours, 400 μg/ml of G418 (345812; Calbiochem EMD, San Diego, CA) was added to select for transfected cells. After 10 days, the cells were collected and lysed, and c-Myc–tagged recombinant mouse NEU3 was purified using a Myc-Trap agarose kit (ytak-20; Chromotek, Hauppauge, NY) following the manufacturer’s protocol. Recombinant protein was checked for protein concentration by OD 260/280/320 using a Take3 micro-volume plate with a SynergyMX plate reader (BioTek, Winooski, VT, USA). NEU3 is 51 kDa, and was further purified by centrifugation at 10,000 xg for 5 minutes at 4°C through an Amicon Ultra 0.5 ml 100 kDa cutoff spin filter (Millipore, Billerica, MA, USA). The NEU3 was analyzed for size and purity by PAGE on, and Coomassie staining of, 4–20% Tris/glycine gels (Bio-Rad, Hercules, CA, USA), as described previously [25, 27-29]. The NEU3 was stored in 50 μl of 10% glycerol, 100 mM glycine, and 25 mM Tris-HCl, pH 7.3, at 4°C.

### Recombinant mouse inactive NEU3 variant generation

Starting with the murine NEU3 expression plasmid MR223297, a variant mutated to change the tyrosines at positions 179 and 181 to phenylalanines was generated using a QuikChange II Site-Directed Mutagenesis Kit (#200523; Agilent, Santa Clara, CA) and the primer 5’ CATCCCCGCC<u>TTC</u>GCC<u>TTC</u>TATGTCTCACGTTGG 3′, with the underscored sequences representing the point mutation sites. Other workers found that these two mutations eliminate NEU3 sialidase activity [18]. The resulting plasmid was sequenced to confirm the point mutations and absence of other mutations.

### Cell Isolation and Culture

Human peripheral blood was collected from healthy volunteers who gave written consent with specific approval from the Texas A&M University human subjects review board. All methods were performed in accordance with the relevant guidelines and regulations. Blood collection, isolation of peripheral blood mononuclear cells (PBMC), and cell culture were done as described previously [22, 23, 25, 30]. Murine spleen cells were isolated by forcing diced spleen fragments through a 100 μm cell strainer (BD Biosciences, San Jose, CA) using the plunger of a 1 ml syringe (BD Medical, Franklin Lakes, NJ), as described previously [31]. To determine the activity of native and mutated NEU3 we assayed the ability of NEU3 to induce extracellular accumulation of IL-6 from human PBMC and murine spleen cells [23, 25], using human (BioLegend) and murine (PeproTech, Cranbury, NJ) IL-6 ELISA kits following the manufacturers’ protocols.

### Mouse models of pulmonary inflammation and fibrosis

This study was carried out in accordance with the recommendations in the Guide for the Care and Use of Laboratory Animals of the National Institutes of Health. The protocol was approved by Texas A&M University Animal Use and Care Committee (IACUC 2020-0272). All procedures were performed under anesthesia, and all efforts were made to minimize suffering. Animals were housed with a 12-hour/12-hour light-dark cycle with free access to food and water, and all procedures were performed between 09:00 and noon. To induce inflammation and fibrosis, 7-8 week old 20-25 g male and female C57BL/6 mice (Jackson Laboratories, Bar Harbor, ME) were given an oropharyngeal aspiration of 3 U/kg bleomycin (2246-10; BioVision Incorporated, Milpitas, CA) in 50 μl of 0.9% saline or oropharyngeal saline alone, as a control, as previously described [22, 23, 25, 32]. Starting 10 days after saline or bleomycin had been administered, some of the mice received an oropharyngeal aspiration of 15 ng of recombinant mouse NEU3 or mutated NEU3 in 50 μl of 0.9% saline, or saline, every 48 hours. An additional cohort of mice received only 15 ng of recombinant mouse NEU3, mutated NEU3, or saline, every 48 hours over the course of 21 days. Three male and three female mice that did not receive saline, bleomycin, or NEU3 were defined as naïve mice. All the mice were monitored twice daily to observe any sign of distress. At the indicated time points, mice were euthanized by CO_2_ inhalation, and bronchoalveolar lavage fluid (BALF) and lung tissue was obtained and analyzed as described previously [22, 23, 25, 32, 33].

### Staining of bronchoalveolar lavage fluid (BALF) cells

BALF cells were counted and processed to prepare cell spots as described previously [22, 23, 25, 32]. After air drying for 48 hours at room temperature, some of the cell spots were fixed and immunochemistry was performed as described previously [22, 23, 25, 32] using anti-CD3 (NB600-1441, rabbit clone SP7, Novus Biologicals, Centennial, CO) to detect T-cells, anti-CD11b (101202, clone M1/70, BioLegend, San Diego, CA) to detect blood and inflammatory macrophages, anti-CD11c (M100-3, clone 223H7, MBL International, Woburn, MA) to detect alveolar macrophages and dendritic cells, anti-CD45 (147702, clone I3/2.3, BioLegend) for total leukocytes, anti-Ly6g (127602, clone 1A8, BioLegend) to detect neutrophils, with isotype-matched irrelevant rat (BioLegend) and rabbit (Novus Biologicals) antibodies as controls. Using a 40x objective, at least 150 cells from each stained BALF spot were examined and the percent positive cells was recorded.

### Lung histology

After collecting BALF, the lungs from the mice were harvested and inflated with Surgipath frozen section compound (#3801480, Leica, Buffalo Grove, IL), frozen on dry ice, and stored at -80°C. 10 μm cryosections of lungs were placed on Superfrost Plus glass slides (VWR) and were air dried for 48 hours. Immunohistochemistry was done as previously described [22, 23, 25, 32] using anti-CD3, anti-CD11b, anti-CD11c, anti-CD45, anti-Ly6g, or anti-Mac2 (clone M3/38; BioLegend) antibodies with isotype-matched irrelevant rat and rabbit as controls. Positively stained cells were counted from randomly selected fields, and presented as the number of positive cells per mm^2,^ as described previously [22, 23, 25, 32]. Lung sections were also stained with Sirius red to detect collagen and analyzed as previously described [23, 34].

### Statistical Analysis

Statistical analysis was performed using Prism v7 (GraphPad Software, La Jolla, CA). Statistical significance between two groups was determined by t test, or between multiple groups using analysis of variance (ANOVA) with Sidak’s post-test, and significance was defined as p<0.05.

## Results

### Recombinant murine NEU3, but not mutated NEU3, has NEU3 activity

We previously observed that the BALF from fibrotic lungs, and fibrotic lesions from human and mouse lungs, have elevated levels of NEU3, that NEU3 inhibitors attenuate bleomycin-induced pulmonary fibrosis, and that *Neu3*^*-/-*^ knockout mice are resistant to bleomycin-induced inflammation and fibrosis [22, 23, 25]. To further elucidate the contribution of NEU3 to fibrosis, we generated recombinant murine NEU3 and a mutated recombinant murine NEU3 protein with two point mutations in the active site. NEU3 and mutated NEU3 were 51 kDa as assessed by PAGE and Coomassie staining (Figure 1A). Recombinant human NEU3 induces the accumulation of IL-6 from human peripheral blood mononuclear cells (PBMC), and this is blocked by NEU3 inhibitors [22, 23, 25], indicating that the effect on IL-6 is dependent on NEU3 activity. The recombinant murine NEU3, but not the mutated NEU3, upregulated extracellular accumulation of IL-6 in both human and murine cells (Figure 1B and C), indicating that the recombinant mouse NEU3 has NEU3 activity and that the mutated NEU3 does not and is thus inactive.

**Figure 1:**
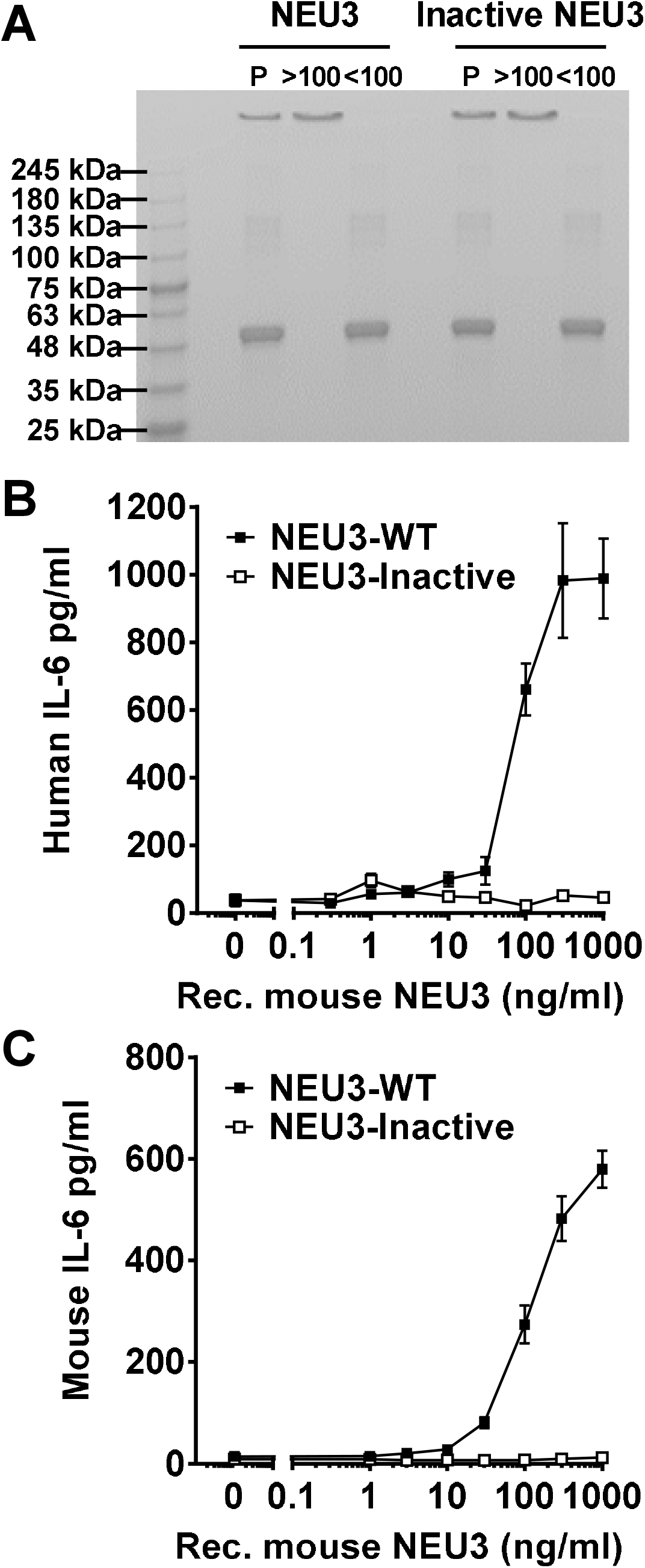
Recombinant murine NEU3, but not inactive NEU3, upregulates IL-6. **A)** Recombinant NEU3 proteins were purified (P) and then filtered through a 100 kDa filter and the retentate (>100) and flow through (<100) were collected. Molecular mass markers in kDa are at left. **B)** Human PBMC and **C)** murine spleen cells were incubated with NEU3 or inactive NEU3 for two days. Supernatants were then assessed for **B)** human or **C)** murine IL-6 by ELISA. Values are mean ± SEM, n = 3.

### Aspiration of NEU3 or inactive NEU3 induces inflammation

To determine if elevated levels of NEU3 protein in the lung, without any other exogenous insult, could induce inflammation and/or fibrosis, mice received 15 ng of either NEU3 or inactive NEU3 every 48 hours by oropharyngeal aspiration. This dose of NEU3 corresponds to the total amount of NEU3 in the BALF of mice at day 21 after bleomycin [23]. There was no discernable effect of NEU3 on the appearance or behavior of the mice, aspiration did not significantly affect body weight, or white fat, liver, kidney, spleen, heart, or brown fat weights as a percent of total body weight (Supplemental Figure 1A-C). Compared to naive controls, NEU3 aspiration increased BAL cell counts and BAL CD11c and CD45 cell counts in male and female mice (Figure 2). Compared to NEU3-treated mice, inactive NEU3-treated mice had lower BAL cell counts (Figure 2A), and this trend was observed in both male and female mice (Figure 2B). Compared to NEU3 aspiration, aspiration of inactive NEU3 caused a smaller increase in CD45 cells in the BAL for female but not male mice (Figure 2C and D). In female mice, NEU3 aspiration decreased the percentage of CD11c cells in the BAL compared to saline or inactive NEU3 aspiration (Figure 2F). There were no significant differences in BAL protein levels between naïve mice (1.2 ± 0.2 mg/ml: mean ± SEM), NEU3 (1.8 ± 0.4 mg/ml; mean ± SEM) and inactive NEU3 (1.4 ± 0.9 mg/ml; mean ± SEM) treated mice.

**Figure 2:**
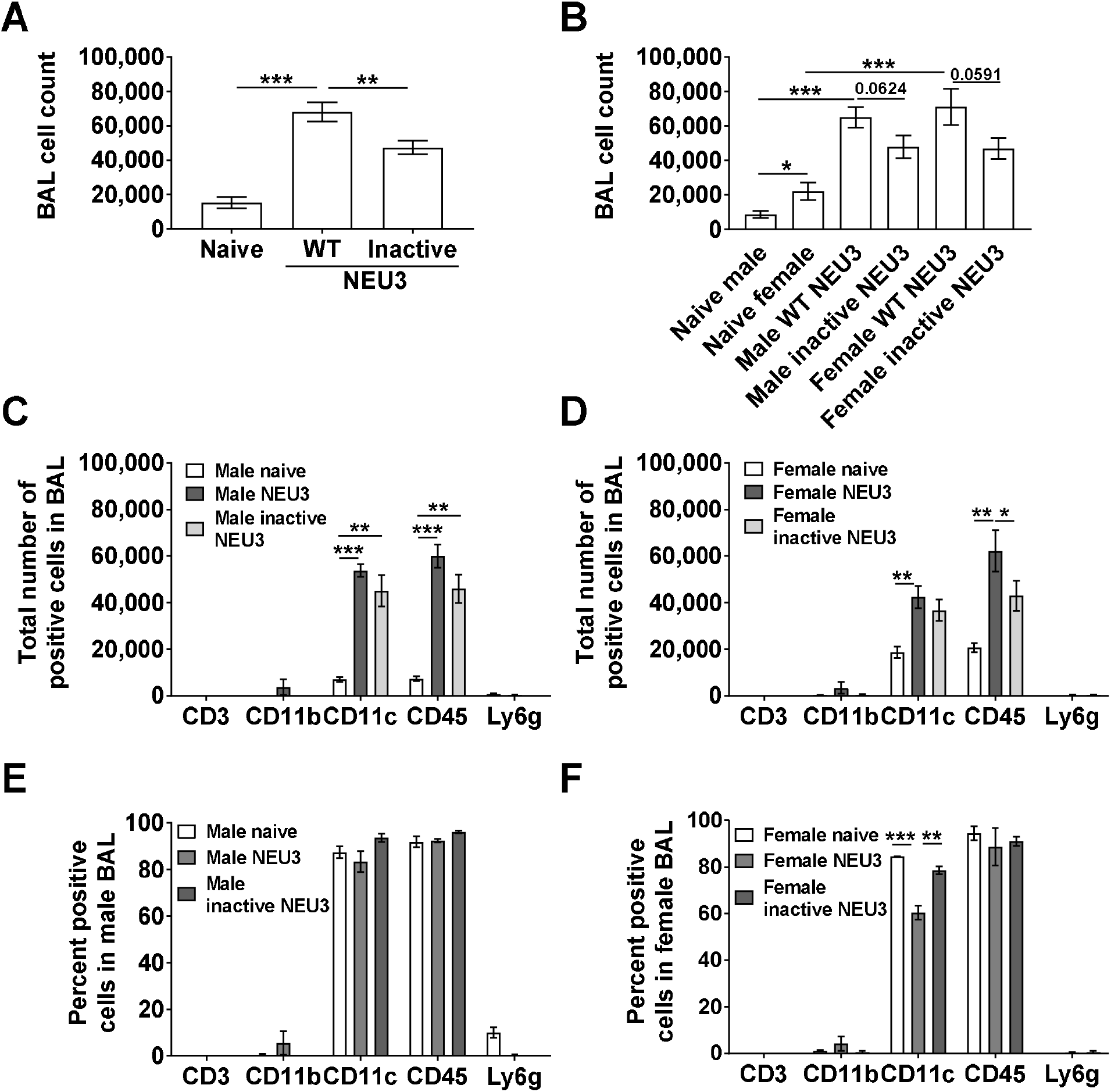
NEU3 treated mice have increased numbers of bronchoalveolar lavage (BAL) cells. **A)** The total number of cells in mouse BAL after the indicated treatment. Values are mean ± SEM, n=6 (3 male and 3 female) mice per group. **B)** BAL cell counts separated into in male and female mice, n = 3 male and 3 female mice per group. **C-F)** BAL cell spots at day 21 were stained for the markers CD3, CD11b, CD11c, CD45, and Ly6g, and the percent of cells stained was determined in five randomly chosen fields of 100–150 cells, and the percentage was multiplied by the total number of BAL cells for that mouse to obtain the total number of BAL cells staining for the marker. Values are mean ± SEM., n = 3. *p < 0.05; **p < 0.01, *** p < 0.001 (one-way ANOVA, Sidak’s test).

After removing BAL, lung sections were stained to detect immune cells that were not removed by lavage. In male mice, compared to naive mice, NEU3 increased the number of CD3, CD11b, CD45, and Mac2 positive cells in the lungs (Figure 3A-F). In female mice, NEU3 decreased CD3 positive cells and increased Mac2 and Ly6g positive cells (Figure 3E and F). Compared to NEU3 aspiration, aspiration of inactive NEU3 did not significantly affect counts of cells expressing any of the 6 markers. Together, the data indicate that aspiration of either form of NEU3 induces inflammation in the lungs of mice.

**Figure 3:**
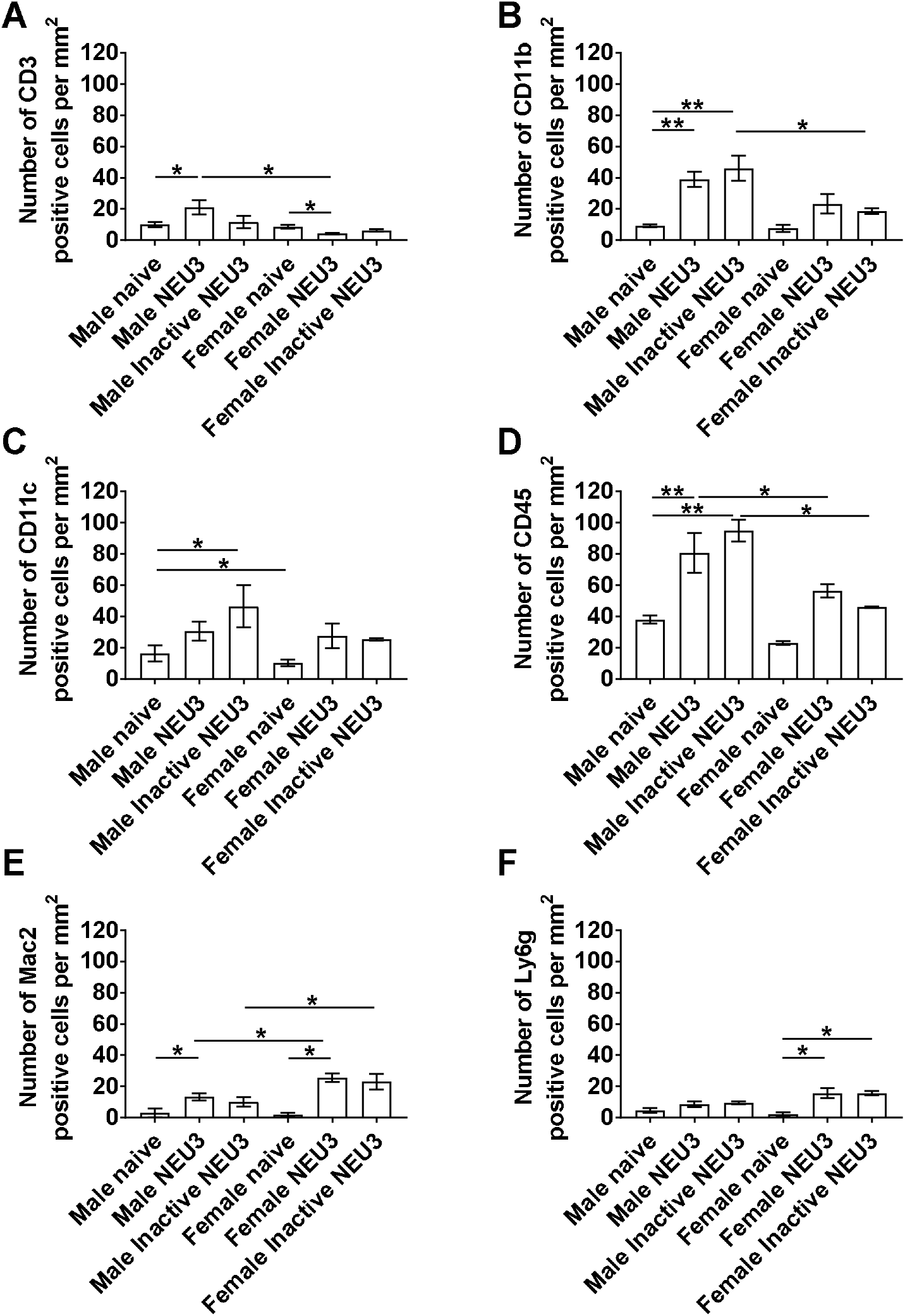
Increase in immune cells in lungs post-BAL of NEU3 treated mice. Cryosections of mouse lungs were stained for **A)** CD3 **B)** CD11b, **C)** CD11c, **D)** CD45, **E)** Mac2 and **F)** Ly6g and counts were converted to the number of positive cells per mm^2^. Values are mean ± SEM, n = 3. *p < 0.05; ** p < 0.01 (one-way ANOVA, Sidak’s test).

### Aspiration of NEU3 but not inactive NEU3 induces fibrosis

Lung sections were also stained with picrosirius red to detect total collagen (Figure 4A-F). In male and female mice, aspiration of NEU3 but not inactive NEU3 caused an increase in picrosirius red staining (Figure 4). These data suggest that NEU3 induces fibrosis, and that the effect is dependent on NEU3 activity.

**Figure 4:**
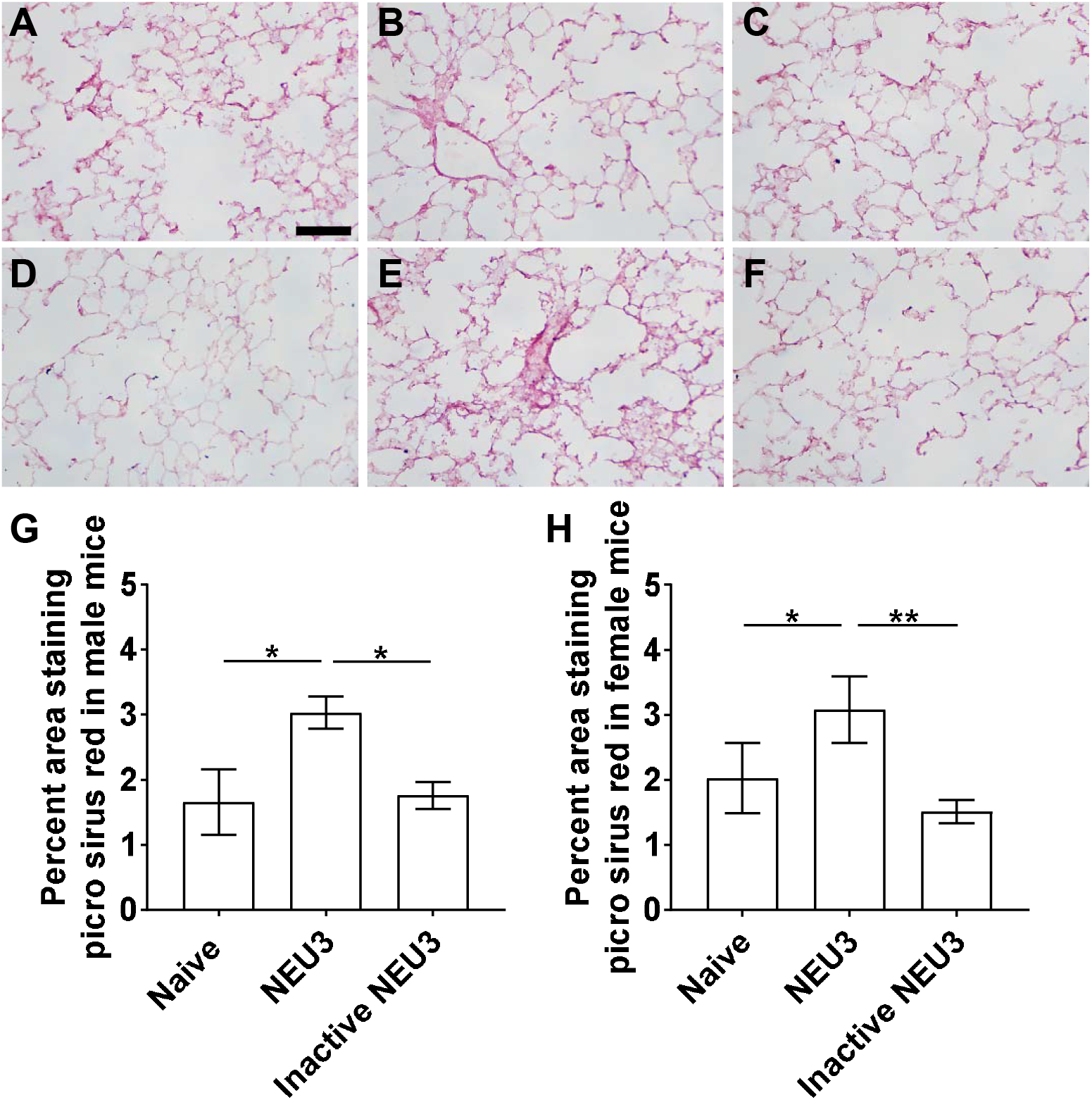
NEU3 but not inactive NEU3 induces fibrosis. Sections of lung tissue from naïve mice and mice treated with NEU3 or inactive NEU3 were stained with picrosirius red to show collagen content. **A)** Naïve, **B)** NEU3, and **C)** inactive NEU3 treated male mice, and **D)** Naïve, **E)** NEU3, and **F)** inactive NEU3 treated female mice. All images are representative of three mice per group. Bar is 0.1 mm. Picrosirius red quantification of **G)** male and **H)** female mice. Values are mean ± SEM, n = 3. *p < 0.05; **p < 0.01 (one-way ANOVA, Sidak’s test).

### Following bleomycin, aspiration of NEU3 but not inactive NEU3 increases inflammation in male mice

Bleomycin aspiration is a standard way to increase inflammation and fibrosis in the lungs of mice [35, 36], and we found that this also increases levels of NEU3 in the lungs [22, 23, 25]. To determine if further increasing levels of NEU3 in the lungs would alter inflammation and fibrosis after aspiration of bleomycin, mice were treated with saline or bleomycin at day 0, and then from days 10 to 20 received 15 ng of either NEU3, inactive NEU3, or saline control every 2 days by oropharyngeal aspiration. As observed previously, male mice that received bleomycin have a drop in body weight between days 3 and 10, but there was no significant weight loss in the female mice (Figures 5A and B). Aspiration of NEU3 or inactive NEU3 did not significantly alter the weights of the mice (Figures 5A and B). As previously observed, compared to saline controls, bleomycin instillation led to an increase in the number of cells recovered from the BALF in both male and female mice at day 21 (Figure 5C and 5D). In the lungs of male mice that received bleomycin, NEU3 aspiration had significantly higher BAL cell counts compared to saline or inactive NEU3 treated mice (Figure 5C). In female mice that received either saline or bleomycin, NEU3 aspiration did not significantly alter BAL cell counts (Figure 5D).

**Figure 5:**
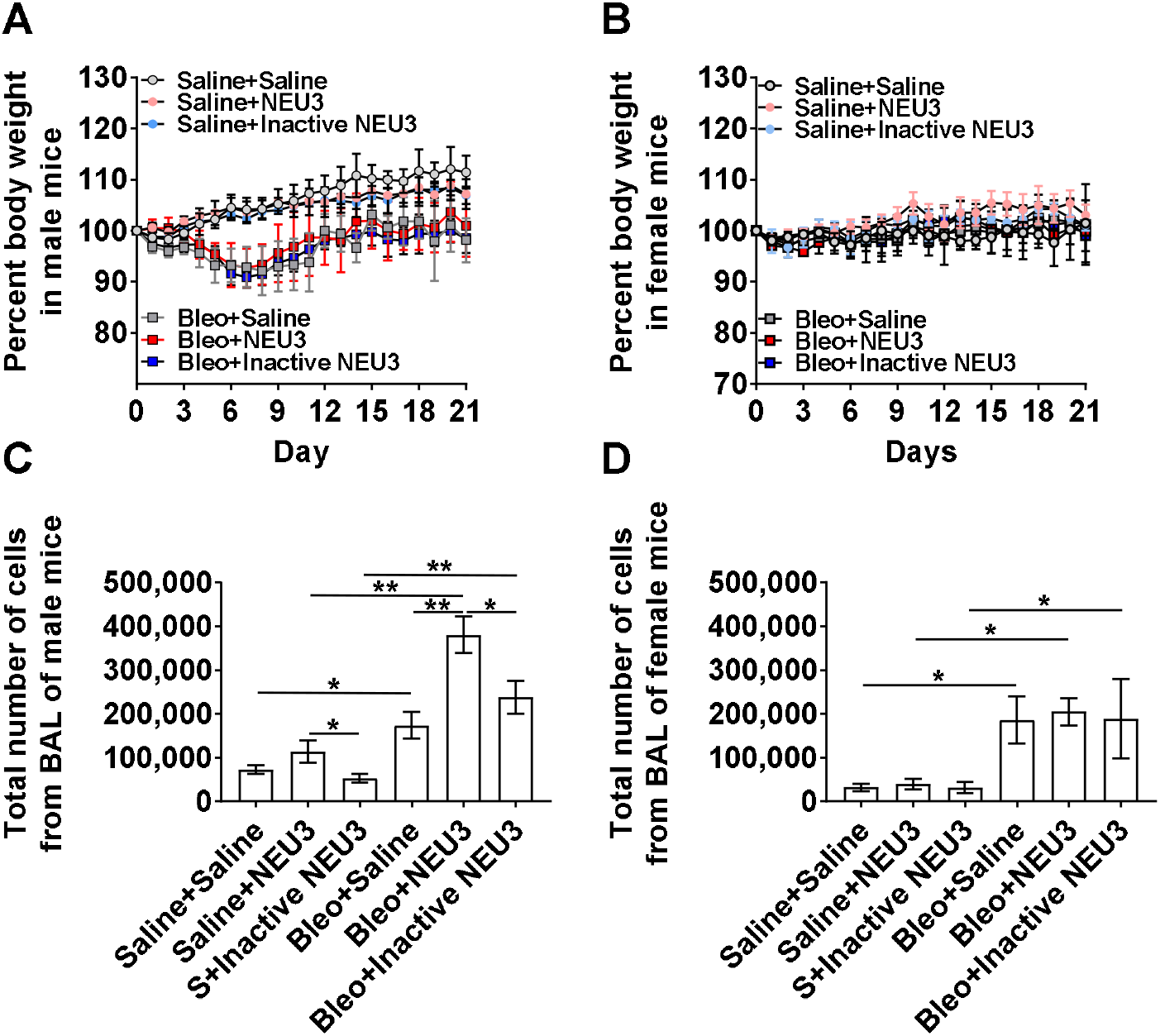
NEU3 increases the number of BAL cells in bleomycin treated male mice. Percent change in body weight in **A)** male and **B)** female mice after saline (S) or bleomycin (Bleo) aspiration at day 0, and saline, NEU3, or inactive NEU3 treatment from days 10 to 20. Values are means ± SEM, n = 3 mice per group. The total number of cells in **C)** male and **D)** female mouse BAL after the indicated treatment. Values are mean ± SEM, n = 3. *p < 0.05; **p < 0.01 (one-way ANOVA, Sidak’s test).

In bleomycin-treated male but not female mice, aspiration of NEU3 but not inactive NEU3 increased the number of CD45 positive cells in the BAL at day 21 (Figures 6A and C). No treatment affected the percentages of CD3, CD11b, CD11c, CD45, or Ly6g positive cells in the BAL for male or female mice (Figures 6B and D). In the lungs of bleomycin-treated male mice after BAL, NEU3 aspiration, but not aspiration of inactive NEU3, increased the numbers of CD11c and CD45 positive cells (Figure 7A). Together, the results suggest that in male but not female mice, increasing levels of NEU3 starting at day 10 after bleomycin causes a further increase in inflammation.

**Figure 6:**
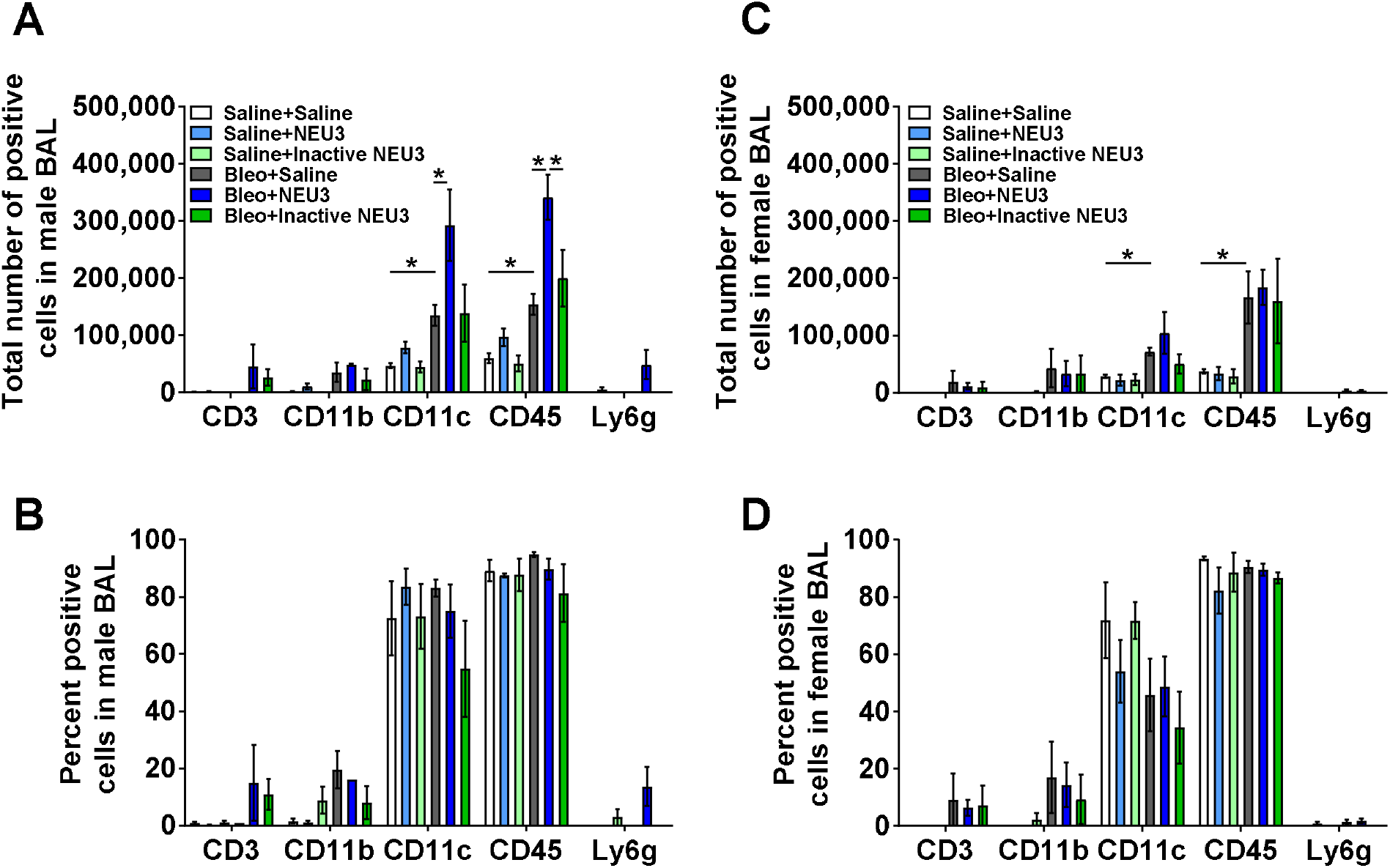
NEU3, but not inactive NEU3, treated male mice have increased numbers of inflammatory immune cells in the BAL following bleomycin aspiration. Graphs show total number of BAL cells in **A)** male and **B)** female mice, and the percent of cells in **C)** male and **D)** female mice, staining for the markers CD3, CD11b, CD11c, CD45, and Ly6g. Values are mean ± SEM, n = 3. *p < 0.05 (one-way ANOVA, Sidak’s test).

**Figure 7:**
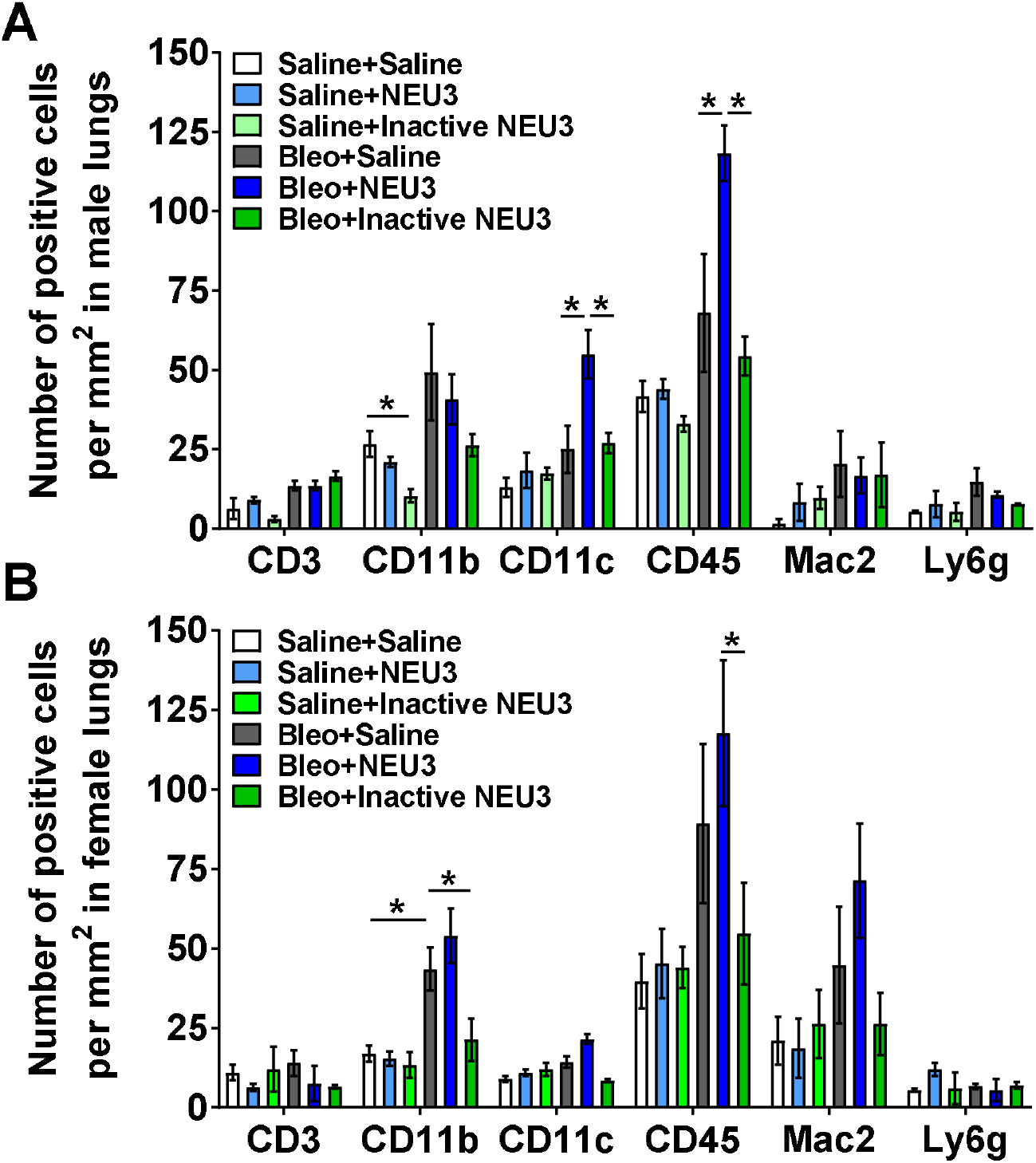
NEU3, but not inactive NEU3, treated male mice have increased numbers of inflammatory cells in lung tissue following bleomycin aspiration. Cryosections of **A)** male and **B)** female mouse lungs were stained for CD3, CD11b, CD11c, CD45, Mac2, and Ly6g, and counts were converted to the number of positive cells per mm^2^. Values are mean ± SEM, n = 3. *p < 0.05; (one-way ANOVA, Sidak’s test).

### Aspiration of NEU3 but not inactive NEU3 for 10 days increases fibrosis

Lung sections were also stained with picrosirius red to detect total collagen. In male and female mice, aspiration of NEU3, but not inactive NEU3, for 10 days increased picrosirius red staining (Figure 8M-N). In bleomycin treated mice, NEU3 aspiration increased picrosirius red staining in male but not female mice (Figure 8M-N). In male and female mice treated with bleomycin, compared to aspiration of NEU3, aspiration of inactive NEU3 reduced picrosirius red staining (Figure 8M-N). These data suggest that increasing NEU3 can increase fibrosis, and that this effect requires NEU3 activity.

**Figure 8:**
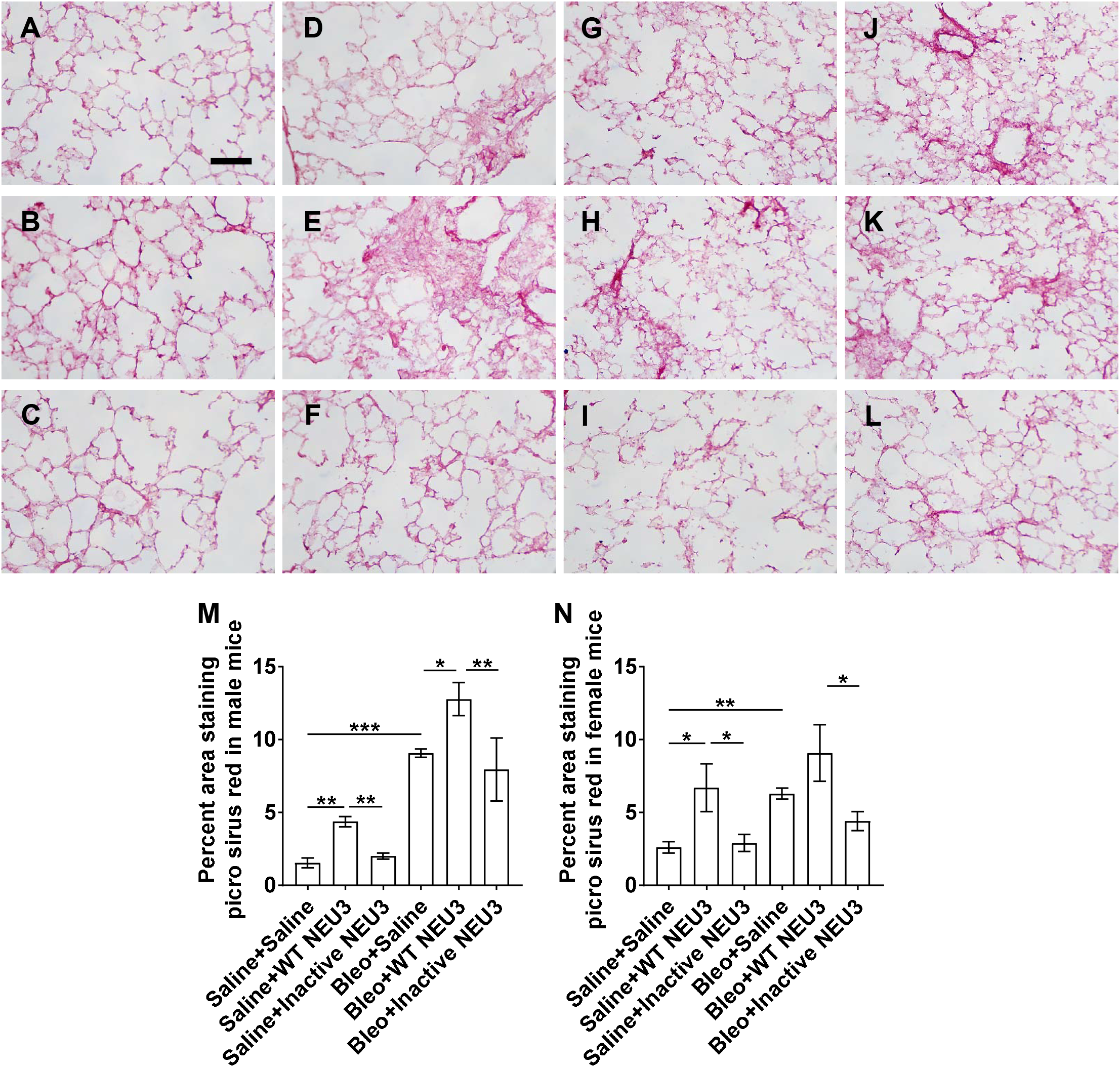
Enhanced fibrotic response in NEU3 treated mice following bleomycin. Sections of lung tissue from mice treated with saline or bleomycin (Bleo) aspiration at day 0, and NEU3 or inactive NEU3 treatment from days 10 to 20, were stained with picrosirius red to show collagen content. **A-F)** Male and **G-L)** female mice aspirated with **A and G)** saline at day 0, **B and H)** saline at day 0 and NEU3 from days 10-20, **C and I)** saline at day 0 and inactive NEU3 from days 10-20, **D and J)** bleomycin at day 0 and saline from days 10 to 20, **E and K)** bleomycin at day 0 and NEU3 from days 10-20, **F and L)** bleomycin at day 0 and inactive NEU3 from days 10-20. All images are representative of three mice per group. Scale bar is 0.1 mm. Picrosirius red quantification of **M)** male and **N)** female mice. Values are mean ± SEM, n = 3. *p < 0.05; **p < 0.01 ***p <0.001 (one-way ANOVA, Sidak’s test).

## Discussion

In this report, we observed that the murine sialidase NEU3, but not enzyme inactive murine NEU3, can induce IL-6 production from human and murine cells as observed previously for human NEU3 [23, 37]. We also observed that NEU3 treatment increased the number of immune cells in the BAL and lung tissue both alone and after instillation of bleomycin, compared to naïve and saline treated mice respectively. Inactive NEU3 did increase the number of cells in the BAL compared to naïve mice, but the response was less than that of enzyme active NEU3. We also observed that in male mice, NEU3 augmented the number of cells in the BAL following bleomycin, and that inactive NEU3 did not augment the inflammatory response compared to active NEU3. NEU3 treatment also upregulated the fibrotic response in both male and female mice when given alone, but only NEU3, and not inactive NEU3, augmented the fibrotic response after bleomycin challenge.

Elevated NEU3 levels are found during inflammation and fibrosis in many organ systems, *Neu3*^*-/-*^ knockout mice show an attenuated inflammatory and fibrotic response, and NEU3 inhibitors reduce these responses in wild type mice [19-26, 38-41]. We observed that the instillation of exogenous active NEU3, but not the inactive NEU3, augmented bleomycin-induced lung inflammation in male mice, and fibrosis in male and female mice. This suggests that the observed effects of NEU3 on inflammation and fibrosis are due to the enzymatic activity of NEU3 rather than the presence of the NEU3 protein itself. Other workers, and this report, observed that compared to male mice, female mice have a reduced fibrotic response to bleomycin ([42, 43] and Figure 8). We observed in this report that female mice have a reduced fibrotic response to aspirated NEU3 following bleomycin. This suggests that compared to male mice, female mice may have a reduced sensitivity to NEU3. Despite these sex differences, our results indicate that NEU3 potentiates pulmonary fibrosis.

## Figure legends

**Supplementary Figure 1.**
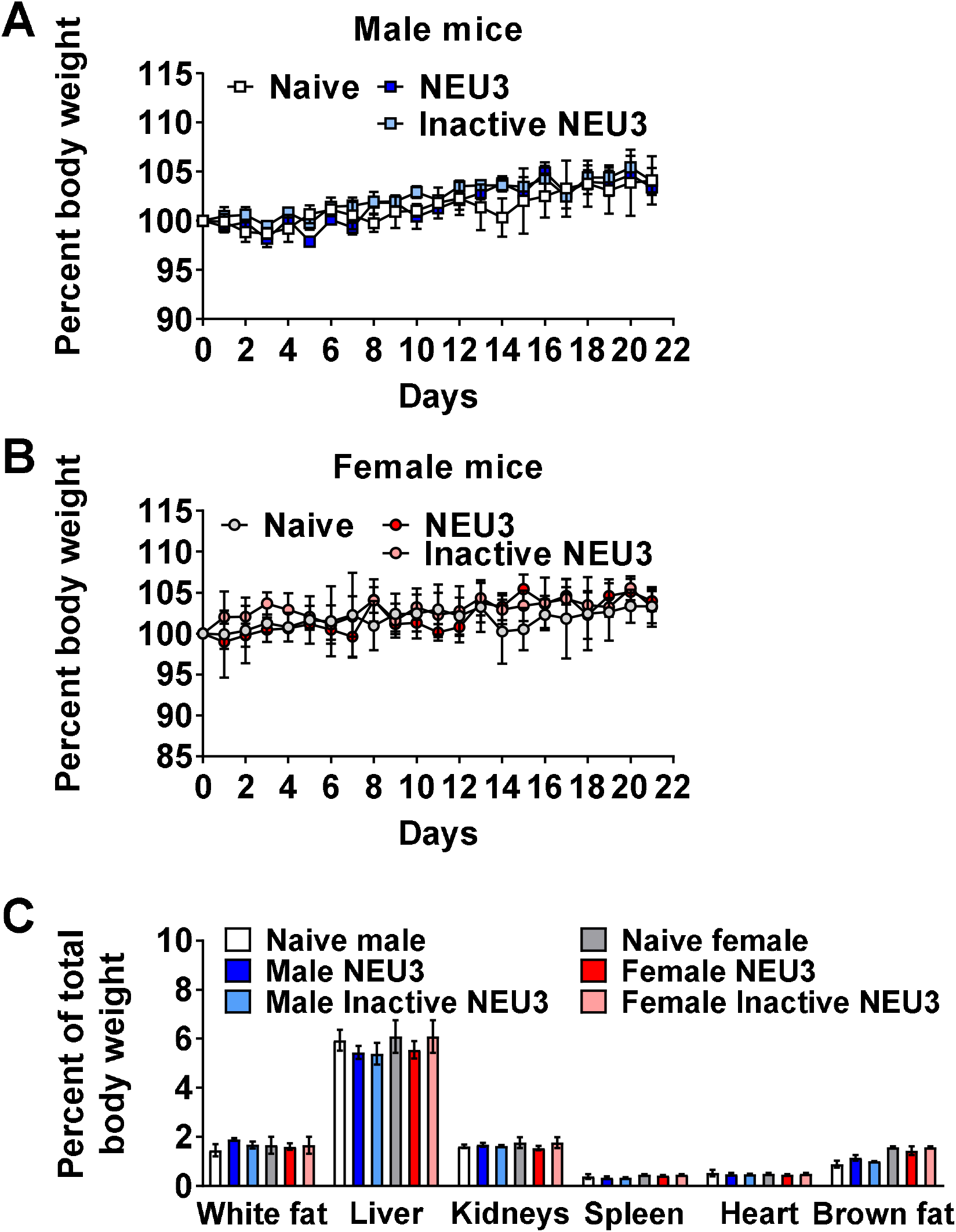
Aspiration of NEU3 or inactive NEU3 had no significant effect on body weight or organ weights. Percent change in body weight of **A)** male and **B)** female naïve mice, or mice after aspiration of NEU3 or inactive NEU3 for every 48 hours for 20 days. **C)** Weights of white fat, liver, kidneys, spleen, heart, and brown fat as percentage of total bodyweight at day 21. Values are mean ± SEM, n = 3. There were no significant changes as determined by t test or one-way ANOVA, Sidak’s test.

